# Modifying adaptive therapy to enhance competitive suppression

**DOI:** 10.1101/2020.10.26.355701

**Authors:** Elsa Hansen, Andrew F. Read

## Abstract

Adaptive therapy is a promising new approach to cancer treatment. It is designed to leverage competition between drug-sensitive and drug-resistant cells in order to suppress resistance and maintain tumor control for longer. Prompted by encouraging results from a recent pilot clinical trial, we evaluate the design of this initial test of adaptive therapy and identify three simple modifications that should improve performance. These modifications are designed to increase competition and are easy to implement. Using the mathematical model that supported the recent adaptive therapy trial, we show that the suggested modifications further delay time to tumor progression and also increase the range of patients who can benefit from adaptive therapy.

## 1. Introduction

Drug discovery and regimen design play complementary roles in advancing cancer therapy. Although drug discovery receives a disproportionate amount of attention [1–3], regimen design is critically important and has been responsible for dramatic advancements in patient care. Regimen design, for example, played a key role in making childhood acute lymphoblastic leukemia (ALL) an essentially curable disease [4–6]. Regimen design often involves identifying synergistic combinations [7–9] and sequences of drugs [10–12]. As discussed here, however, improving regimen design does not have to involve changing or adding therapeutics. It is possible to enhance patient outcomes simply by modifying how the current drugs are used.

We consider an innovative treatment regimen that is showing very promising results in a pilot clinical trial [13]. This novel regimen, called “adaptive therapy”, uses the same drug as the standard of care, but applies it differently. This emphasizes that simply changing how we use therapeutics can lead to dramatic improvements in patient care. Although these preliminary results must be interpreted with caution, they suggest that – in certain circumstances – adaptive therapy may represent a promising new paradigm for patient treatment.

Adaptive therapy has two important but distinct features. First, it is “adaptive” [14]. Instead of using a fixed schedule, treatment decisions are based on how individual tumors respond to treatment. This is in contrast to the majority of regimens which use predetermined treatment schedules. Second, it is designed to leverage competition between drug-sensitive and drug-resistant cells to improve tumor control (Figure 1). This differs from the common goal of achieving and/or sustaining large tumor responses. Rather than attempting to drive tumor cell populations to undetectable levels, it deliberately maintains a notable tumor burden in order to competitively suppress resistance [15]. Competitive suppression has been shown to work in theory and experiment (both in vitro and in vivo) for both cancer and infections [13,16–24]. The recent trial of adaptive therapy provides evidence that an easily implementable realization of competitive suppression may work in the clinic [13].

**Figure 1.**
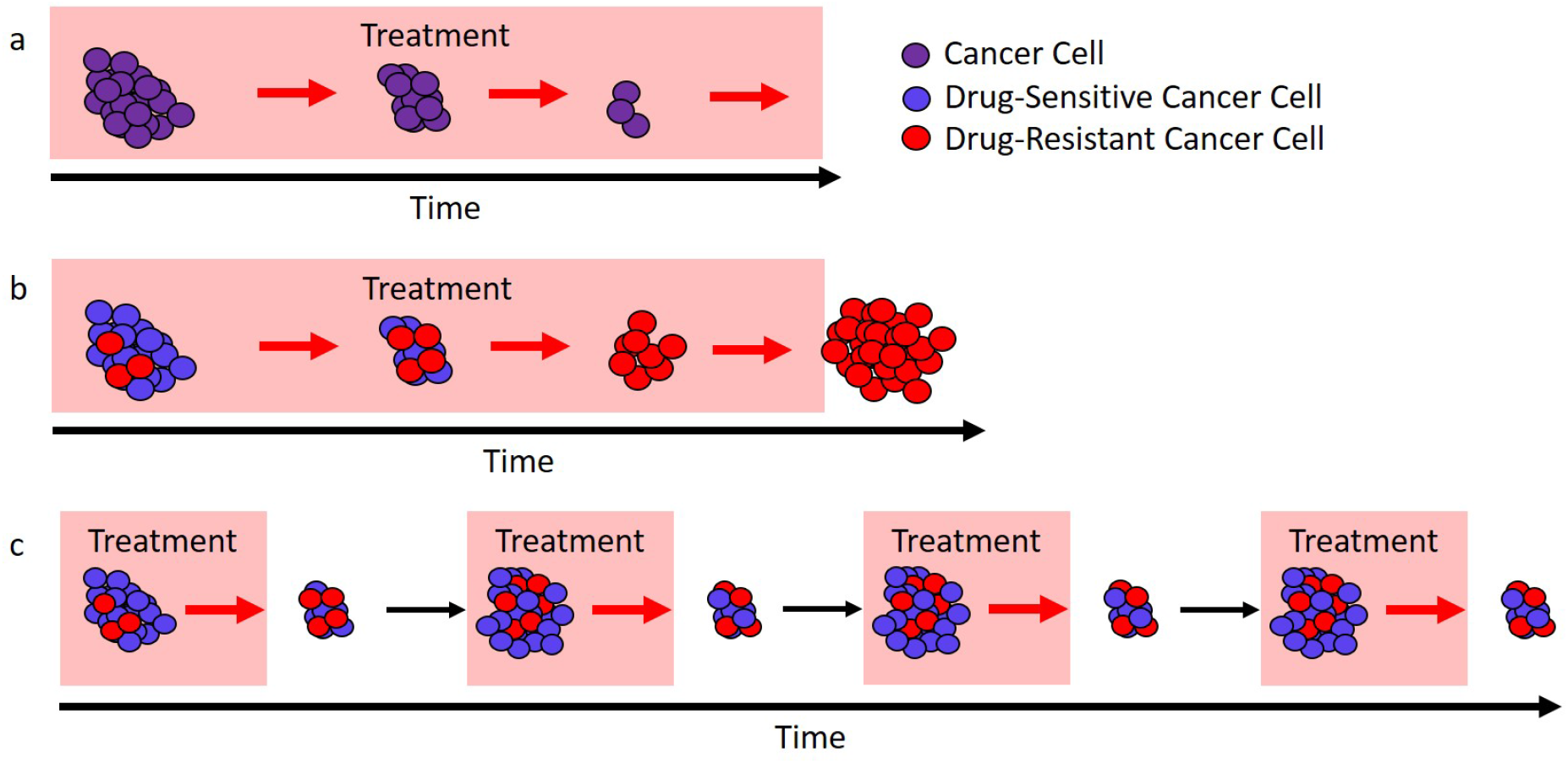
Different approaches to treatment. (**a**) In ideal circumstances continuous treatment (here called the “standard of care”) will stop tumor cell growth and lead to cure. (**b**) If there are drug-resistant cells (red), the standard of care can only remove the drug-sensitive cells (blue) and this leaves a fully resistant tumor that can expand in size. (**c**) If treatment is modulated so that not all drug-sensitive cells (blue) are removed then these cells can competitively suppress the expansion of the resistant cells (red). In ideal circumstances this would lead to longer tumor control.

Although this particular trial tested adaptive therapy on metastatic castrate resistant prostate cancer (mCRPC) [25,26], we are interested in adaptive therapy as a general approach to cancer treatment. Recent theoretical studies have considered the enhancement of adaptive therapy by using multiple drugs [27,28]. Here we evaluate the design of adaptive therapy within the scope of a single drug and discuss how simple modifications to the design should improve patient outcomes. The modifications we suggest are not specific to prostate cancer but are intended to enhance competition in general. This analysis rests on two important assumptions. First we take as a basic tenet that “larger populations generate more competition”. In essence, drug-sensitive cells should competitively suppress the expansion of the resistant population and the larger the sensitive population the stronger this effect should be. Second, we assume that the success of adaptive therapy can be attributed to competitive suppression (see Box 1).

### Box 1. Is competition the reason adaptive therapy works?

The biology of prostate cancer and treatment is complicated. Even though adaptive therapy is designed to leverage competition, it is quite possible that it works for reasons other than competition. For example, improved outcomes may be due to adaptation to different local environments (similar to the hypothesized mechanism of success for bipolar androgen therapy [29,30]). If success is due to some factor other than competition, changes intended to enhance competition may actually reduce the effectiveness of adaptive therapy. This means that modifications should be made with care. Gaining a better understanding of what is causing the current success of adaptive therapy is critical if we are to (i) optimize regimen design and (ii) successfully apply adaptive therapy to other cancers.

## 2. Results

### 2.1. A primer on adaptive therapy

Adaptive therapy maintains a measurable (but contained) tumor burden by modulating treatment according to tumor response. There are a variety of ways that this can be implemented [13,20]. Here we focus on the specific approach taken in a recent pilot clinical trial for mCRPC (NCT02415621) [13]. In this trial, prostate specific antigen (PSA) [31] is used as a proxy for tumor burden and treatment is administered according to the “50% rule” (Figure 2a).

**Figure 2.**
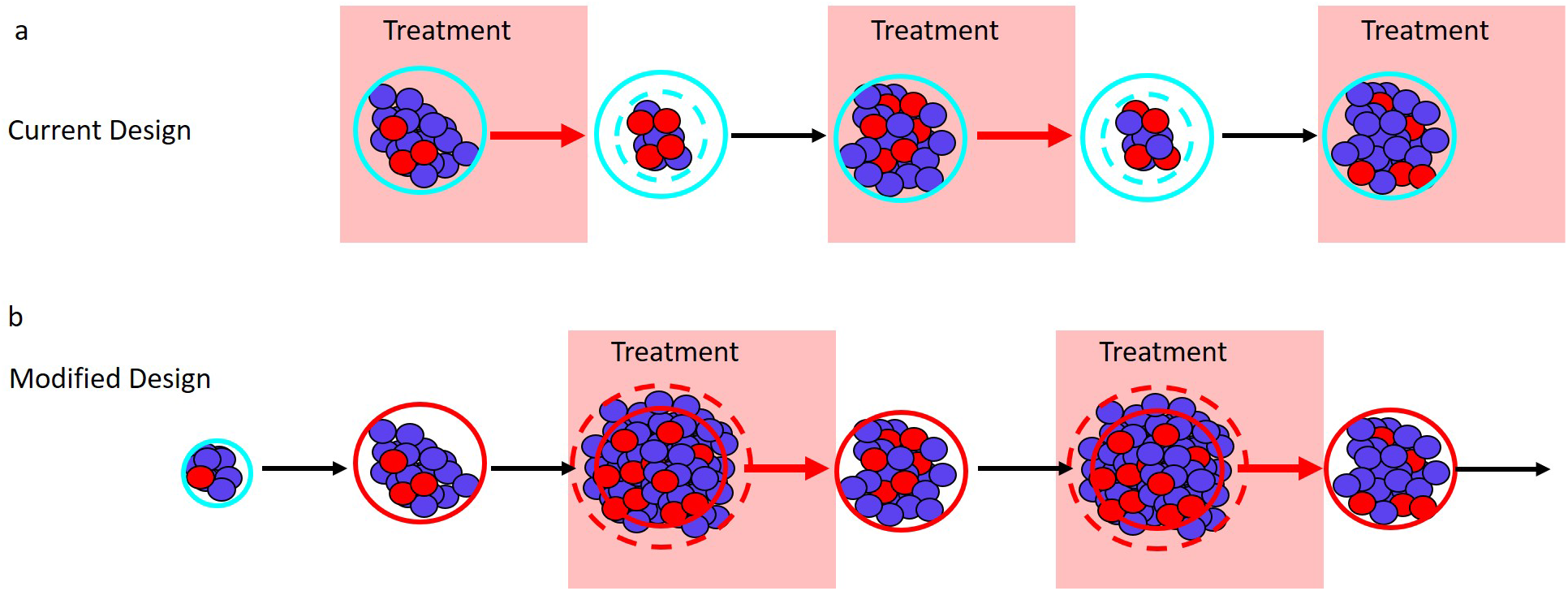
Adaptive therapy designs. PSA is used as a proxy for tumor burden. (**a**) Current adaptive therapy design. Treatment decisions are based on the “50% rule” using the patient’s initial pretreatment baseline PSA (solid blue circle). Treatment is stopped when the PSA falls to 50% of the baseline PSA (dashed blue circle) and is re-initiated only once the PSA returns to its initial baseline. (**b**) Modified regimen design. If the initial baseline PSA (blue circle) is low the PSA is allowed to increase to a larger acceptable baseline PSA (solid red circle). Treatment begins when the PSA exceeds the acceptable baseline by a measurable amount (dashed red circle) and stops once the PSA returns to the acceptable baseline (solid red circle).

Before adaptive therapy begins, a patient’s PSA is determined to establish a baseline from which all treatment decisions will be made. Treatment is then administered until PSA is reduced to 50% of initial baseline. Once a 50% reduction has been observed, treatment is halted and withheld until PSA returns to the initial baseline. This completes a single cycle of adaptive therapy and cycles are repeated until treatment can no longer prevent the PSA from exceeding the initial baseline (i.e., until PSA progression).

This approach differs markedly from the standard of care which treats continuously until progression. Because adaptive therapy halts treatment every time a 50% reduction in PSA is achieved, the expectation is that an appreciable drug-sensitive population is maintained and that this population will slow the expansion of the drug-resistant population — leading to prolonged tumor control (Figure 1). Preliminary results are extremely promising with median time to progression of at least 27 months [13] compared to a median of 9 months (for PSA progression) and 14 months (for radiographic progression) for a contemporaneous cohort which received the standard of care [13].

### 2.2. The role of tumor size and resistance frequency in adaptive therapy

A curious feature of this initial test of adaptive therapy is that *absolute* tumor size plays no role in the current design. This is at odds with the general notion that “larger populations generate more competition”. Instead, the “50% rule” bases treatment decisions on the tumor size *relative* to the initial baseline. An additional consequence of the “50% rule” is that resistance frequency determines which patients have the opportunity to benefit from competitive suppression (as described below). Here we describe how small modifications to adaptive therapy can make absolute tumor size central to the design and greatly enhance competitive suppression. Our suggested modifications are easily implementable and preserve the “adaptive” nature of the treatment regimen. Under these modifications, treatment would occur only when the tumor exceeds a predetermined acceptable burden (Figure 2b).

- **Current design: Maximum tumor size is determined by a patient’s initial baseline burden.** Different patients will have different initial burdens when they present for treatment. Despite this, the current adaptive therapy regimen always implements the “50% rule” from the patient’s “initial baseline burden”. This limits the maximum size of the tumor and so lowers the amount of competitive suppression. **Design Modification: Can the patient’s tumor burden be safely increased?** If yes, then the initiation of adaptive therapy should be delayed until this new larger “acceptable baseline burden” is reached. Withholding treatment until the tumor has grown to a larger size should increase the amount of competition and enhance the performance of adaptive therapy. Whether it is acceptable to allow the tumor to grow before initiating treatment will depend on the specific details of the patient and the cancer as well as the size of the initial baseline burden. In general, making this decision will require balancing the possible benefits (e.g., prolonged time to progression, reduced drug use) with the possible risks (e.g., increased metastasis, greater morbidity). The relationship between tumor size and these other factors is not straightforward [32–36]. In the original trial, however, the initial baseline PSAs ranged from 2.42 to 109.4 ng/ml, suggesting that there is a wide range of acceptable PSA levels [13].
- **Current design: The “50% rule” reduces the average tumor size.** In the current design, treatment begins whenever the tumor reaches the baseline burden and stops whenever it falls below 50 percent of the baseline burden. These successive 50% reductions in tumor burden reduces the average size of the population that is generating competition. **Design Modification: What should trigger treatment?** Since larger populations generate more competition, we suggest “inverting” what triggers treatment starts and stops. Treatment should start whenever the tumor burden exceeds the baseline level by a measurable amount (e.g., 10% larger than the baseline burden) and treatment should stop whenever the burden returns to the baseline. Figure 2 shows how this modification shifts the timing of treatment (shaded blocks in panel b are shifted relative to shaded blocks in panel a). This should increase the average size of the population and enhance competition.
- **Current design: Patients with a high resistance frequency cannot benefit from adaptive therapy.** If a patient’s initial resistance frequency exceeds 50%, they will not be able to achieve a 50% reduction in PSA during the first cycle of adaptive therapy. For these patients treatment resembles the standard of care and they are unable to benefit from adaptive therapy. The above suggested modifications of (i) increasing the baseline (whenever acceptable) and (ii) treating only when the burden exceeds this baseline, help to ameliorate this shortcoming. With these modifications, only patients who begin with almost completely resistant tumors will be unable to complete multiple rounds of adaptive therapy. **Design Modification: Is the patient’s initial resistance frequency likely to be low?** Although the previous modifications should allow patients with high resistance frequencies to benefit from adaptive therapy, special consideration should also be given to patients with very low resistance frequencies. Patients with low initial resistance frequencies may do better with the standard of care than with adaptive therapy (Box 2). For this reason, an effort should be made to identify and exclude patients with very low resistance frequencies. This may be difficult to do, but an evaluation of patient treatment history could help.

#### Box 2. How does resistance frequency impact the performance of adaptive therapy?

Resistance frequency impacts the performance of adaptive therapy in two distinct ways.

**Absolute performance:** Lowering the resistance frequency improves the absolute performance of adaptive therapy. This is not surprising since the lower the initial resistance frequency, the longer it will take for resistance to dominate and exceed the baseline burden. This observation does not rely on competitive suppression and the same relationship also holds for the standard of care [37]. Both adaptive therapy and the standard of care perform better when the initial resistance frequency is low.
**Performance relative to the standard of care:** At very low resistance frequencies adaptive therapy may not perform as well as the standard of care. Consider, for example, a completely sensitive tumor (i.e., resistance frequency of zero). In this case, the standard of care may even clear the tumor and result in cure. On the other hand, with adaptive therapy there is an increased risk that resistance will be introduced de novo (e.g., via mutation or epigenetic changes [38–41]). If resistance is successfully introduced then adaptive therapy will control the tumor only until resistance expands to dominate the cancer. Extrapolating from this simple example suggests that if the resistance frequency is sufficiently small, standard of care may be a better treatment option than adaptive therapy. This is precisely what theory predicts [23]. According to theory, the resistance frequency must be sufficiently large before adaptive therapy is the preferred treatment option. Combining the roles of resistance frequency in the absolute and relative performance of adaptive therapy leads to the following rule: “The resistance frequency must exceed a minimum threshold, but beyond that, lower resistance frequencies are better”. This ensures that adaptive therapy is the preferred option while still maintaining the absolute performance of adaptive therapy. It is possible that this minimum threshold is so low as to be inconsequential (i.e., for tumors of interest, resistance frequencies might naturally exceed this minimum level)[42,43]. If this is the case, then resistance frequency need not play a role in adaptive therapy design. The essential observation, however, is that *if* resistance frequency is important, its role should be the opposite of what it is in the current adaptive therapy design. A low resistance frequency – not a high resistance frequency – should preclude the use of adaptive therapy.

Accurately predicting the impact of these design modifications is difficult because the biology of prostate cancer is complicated and poorly understood. For example, it is known that prostate cancer can be heterogeneous with cell populations that vary in their (i) dependence on and (ii) ability to produce androgens [44–46]. How these different cell populations interact and compete may be more complicated than the simple “larger populations generate more competition” tenet that we use. Therefore, as an initial validation step, we have assessed the impact of these modifications using a mathematical model that accounts for these interactions. This model is the same model that was used to design the current adaptive therapy regimen. According to this model, the design modifications we suggest will improve performance (Figure 3). This emphasizes that, at least for the biological understanding used to design the current regimen, the suggested modified regimen will more effectively prolong time to progression.

**Figure 3.**
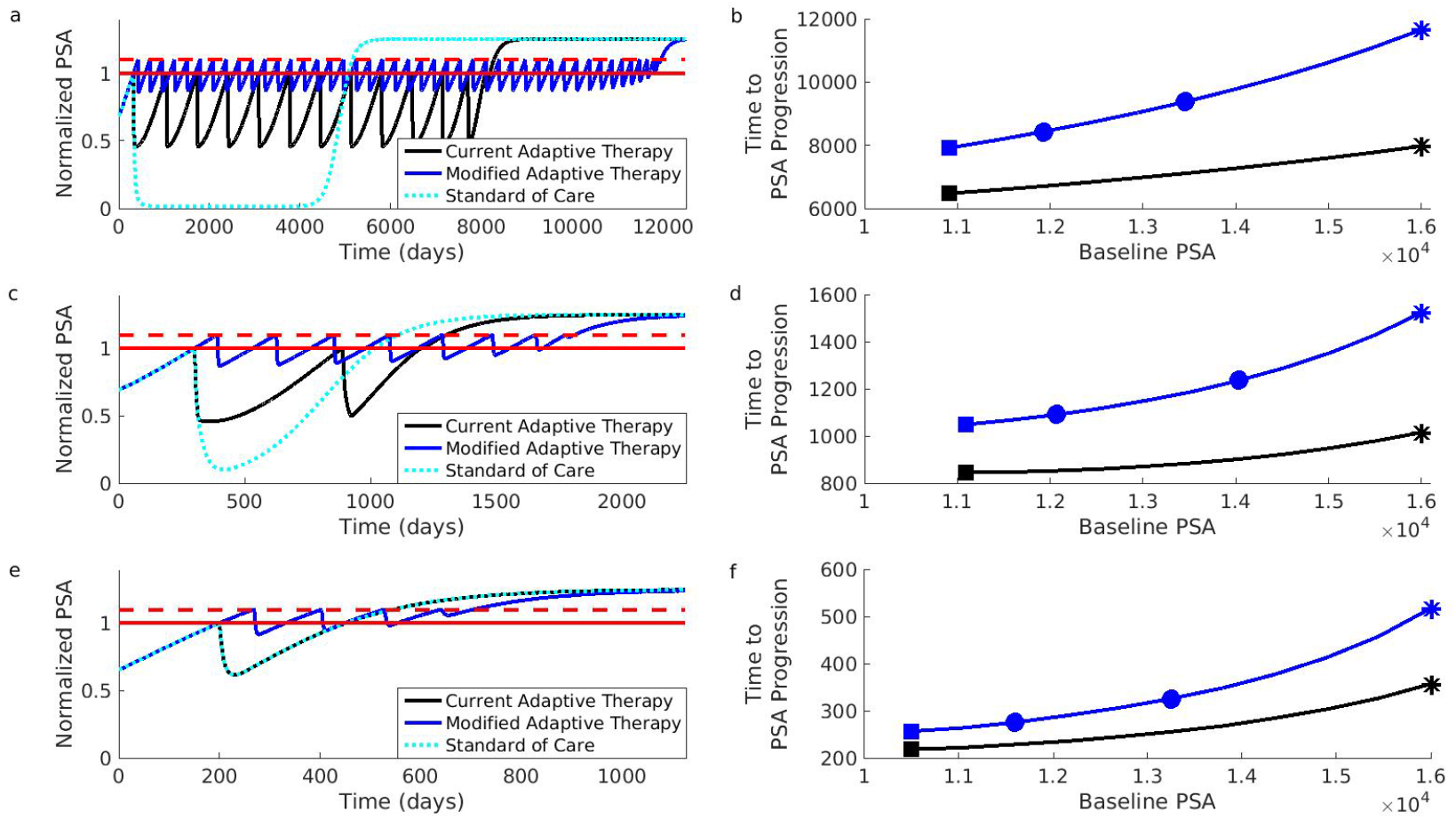
Simulations comparing current and modified adaptive therapy designs. Comparison was performed for three different types of patients. These patients differ in their initial resistance frequencies: low (Patient 1, panels a-b), medium (Patient 2, panels c-d) and high (Patient 3, panels e-r). (**a**) Comparison for Patient 1 assuming that the initial baseline (solid red) cannot be increased. Modified design (blue) controls PSA for longer than the current design (black). Both adaptive therapy designs are better than standard of care (dotted light blue curve). To facilitate comparison, PSA progression for both designs occurs when PSA exceeds 10% of the baseline (dashed red). PSA is normalized relative to initial baseline PSA. (**b**) Time to PSA progression for Patient 1 under the modified (blue) and current (black) adaptive therapy design. If the current design is implemented at the initial baseline indicated by the black square, the modified design implemented at the same baseline would improve results (blue square). But increasing the baseline by 10% (leftmost blue circle) or 25% (rightmost blue circle) improve results further. Asterisks correspond to dynamics shown in (a). (**c-d**) Same as (a-b) except for Patient 2. (**e-f**) Same as (a-b) except for Patient 3. Notice that Patient 3 does not achieve a 50% reduction in PSA during the first cycle of adaptive therapy (black curve in panel (e) is always above 0.5). This means that the current adaptive therapy design coincides with the standard of care (black curve and dotted light blue curve in panel (e) lie on top of each other). Simulations use same mathematical model as the original simulations that supported the design of the current adaptive therapy regimen [13,47]. See Section 4 for details on patient types and simulations.

The original simulations used to develop the current adaptive therapy regimen [13] focused on two types of patients. These patient types differ in how the different cell populations compete and this leads to different initial resistance frequencies. The first patient type (represented by “Patient 1” in Figure 3) exhibits competition dynamics that make resistant cells initially rare. The second patient type (represented by “Patient 2”) is characterized by competition dynamics that promote moderate pretreatment levels of resistant cells. We have also considered a third patient type (“Patient 3”) with an initially high resistance frequency. This third patient type demonstrates that patients can benefit from the modified regimen even if they would have been excluded from the current adaptive therapy trial. (See Section 4 for additional details on patient types.)

Figure 3 shows that the suggested modifications should improve the performance of adaptive therapy for all three types of patients. Even if adaptive therapy must be implemented from a patient’s initial baseline, the modified design results in significant improvements (compare black curves to blue curves). Additionally, these improvements will be enhanced if a larger acceptable baseline can be used. For example, if the current adaptive therapy design results in the performance indicated by the black square (panel b) then the modified design (implemented at the initial baseline) will increase the length of tumor control (blue square, panel b). This improvement can be further enhanced if the baseline can be increased even by modest amounts (leftmost blue square shows improvement for a 10% increase in the baseline, rightmost blue square for a 25% increase, panel b). Further increases will result in further improvements (blue curve is increasing).

## 3. Discussion

Adaptive therapy is a novel treatment paradigm that has shown very promising results in a small clinical trial for mCRPC. Its novelty partly lies in that it is designed to competitively suppress resistance as opposed to minimize tumor burden. Here we have outlined simple modifications that should enhance the performance of adaptive therapy.

According to the tenet “larger populations generate more competition”, our modified design should increase the amount of competitive suppression by (i) increasing the baseline tumor burden whenever possible and (ii) only treating when the tumor exceeds the baseline. An additional benefit of the modified design is that it does not exclude patients with resistance frequencies greater than 50%. This means that more patients have the opportunity to benefit from adaptive therapy.

The main risk of using adaptive therapy is that some patients may do worse than if they had received the standard of care [23]. This is most likely to occur when there is little to no resistance. To mitigate this risk, emphasis should be placed on identifying and excluding patients who are unlikely to harbor much resistance. Here we have not explicitly considered the possibility of cure or additional dynamics which could make adaptive therapy perform worse than the standard of care (e.g., mutation or epigenetic changes [38–41]). In general, these may be possibilities and will complicate the choice between the standard of care and adaptive therapy [48].

In addition to our assumption that larger tumor cell populations generate more competition and adaptive therapy works because of competition (see Box 1) there are two further assumptions underpinning our analysis: (i) PSA is a good proxy for tumor burden [49] and (ii) when PSA progression occurs it can actually be linked to resistance [50]. The comparison we presented uses the same mathematical model that supported the original development of the current adaptive therapy regimen. This model makes very specific assumptions about how cell populations interact (see Section 4). As knowledge of cancer biology and treatment improves, there should be a continuing effort to account for all interactions that will influence performance and inform regimen design.

Our discussion uses mCRPC as an illustrative example because this was adaptive therapy’s first clinical application. We are, however, interested in adaptive therapy as a general approach to cancer treatment. It is essential to emphasize that the performance of adaptive therapy will depend on the cancer, drug and patient population. The modifications we suggest are not specific to prostate cancer but are intended to enhance competition in general. We are certainly not claiming that our design modifications lead to the optimal design. But they are simple, easily implementable modifications that maintain the adaptive nature of adaptive therapy. Regardless of why adaptive therapy is working, or whether it can be improved – the results from the current clinical trial make one thing clear. Having effective therapeutics is not enough; how you use them matters.

## 4. Materials and Methods

### 4.1. Mathematical Model

Prostate cancer cells often rely on androgens to grow. Treatment normally aims to exploit this dependence by employing some form of androgen deprivation therapy (ADT). Even though ADT is initially successful, resistance almost inevitably develops and the cancer progresses. There are a variety of mechanisms that can contribute to this resistance [40,51,52] but variation in androgen dependence is often a contributing factor. The mathematical model used to conceptualize the current adaptive therapy design assumes that certain cells are less dependent on androgens [13,47]. These cells (called androgen independent and denoted by *I*) are resistant to ADT and are responsible for tumor progression during treatment. We use the same model here to compare the current and modified adaptive therapy designs.

This model involves three different populations of cancer cells: (i) androgen dependent (*D*), (ii) androgen independent (*I*) and (iii) androgen producing (*P*). The mathematical equations describing the interactions of these populations are:

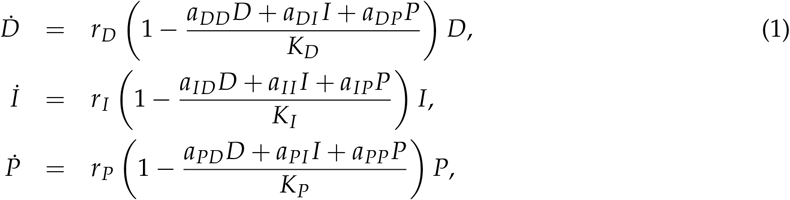

where (i) *r_D_, r_I_* and *r_P_* are the per capita growth rates of the respective populations in the absence of competition, (ii) *K_D_, K_I_* and *K_P_* are the carrying capacities of the respective populations and (iii) the *aij* are constants describing how competition between populations impacts growth.

According to these equations the different populations exhibit logistic growth with both intra- and inter-population competition. Consider, for example, the dynamics of the androgen dependent population (D) described by Equation (1). In the absence of any competition, growth would simply be proportional to the size of the population (i.e., Equation (1) would become *D* = *r_D_D*). Intra-population competition, however, reduces this growth by the factor 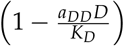. Accounting for competition from all populations reduces growth even further (i.e., by the factor 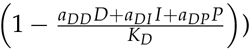.

An essential feature of this model is that the different cell populations interact with each other in two ways. The first is competition (as described through the coefficients *a_ij_*). Competition reduces the growth capacity of each population. The second interaction occurs between the androgen dependent (*D*) and androgen producing cells (*P*). Increasing the androgen producing population (*P*) increases the amount of available androgen and this increases the carrying capacity of the androgen dependent cells (*D*). Mathematically this is encoded by making the carrying capacity of the androgen dependent population (*K_D_*) proportional to the size of the androgen producing population (i.e., *K_D_* = *αP*). Increasing *K_D_* reduces the effect of competition (from all populations) on the androgen dependent population (D). Other types of interactions are not included in the model. For example, cells cannot move between populations through processes like mutation or epigenetic changes [38–41].

This model assumes that treatment modulates dynamics by changing the carrying capacities of the androgen dependent population (*K_D_*) and the androgen producing population (*K_P_*). When there is no treatment these carrying capacities are *K_D_* = 1.5*P* and *K_P_* = *K_I_*. During treatment these carrying capacities are reduced to *K_D_* = 0.5*P* and *K_P_* = 0.01*K_I_*. Treatment has no direct effect on the androgen independent population (I).

### 4.2. Parameter Values and Simulation Details

The parameter values used to produce Figure 3 are the same as the ones used in [13]. The intrinsic growth parameters are 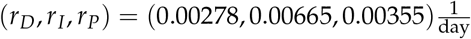 and the carrying capacity for the androgen independent population is *K_I_* = 10000. Two different sets of competition coefficients are used to simulate the dynamics of the two patient types described in [13]. We have also included a third patient type which does not respond well to the current adaptive therapy design. Coefficients for this third patient type were chosen to be identical to the “non-responder” patient type described by Cunningham et al. in an extended analysis of the original mathematical model [47]. The competition coefficients for the different patient types are given in Table 1.

**Table 1.** Competition coefficients for different patient types

| | $a_{DD}$ | $a_{DI}$ | $a_{DP}$ | $a_{ID}$ | $a_{II}$ | $a_{IP}$ | $a_{PD}$ | $a_{PI}$ | $a_{PP}$ |
| --- | --- | --- | --- | --- | --- | --- | --- | --- | --- |
| Patient 1 | 1 | 0.8 | 0.7 | 0.6 | 1 | 0.9 | 0.4 | 0.5 | 1 |
| Patient 2 | 1 | 0.8 | 0.6 | 0.5 | 1 | 0.9 | 0.4 | 0.7 | 1 |
| Patient 3 | 1 | 0.9 | 0.7 | 0.5 | 1 | 0.8 | 0.4 | 0.6 | 1 |

Changes in PSA are described by 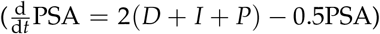. Initial conditions for the simulations were determined in the same way as in [13]. Cell populations were initialized to the values shown in Table 2. The populations were then allowed to grow in the absence of treatment until the PSA reached the desired baseline. Once the PSA reached the desired baseline, adaptive therapy (either the current or modified version) was initiated. The range of PSA baselines used in Figure 3 was 2[*D*_0_ + *I*_0_ + *P*_0_, 0.8*K_I_*].

**Table 2.**
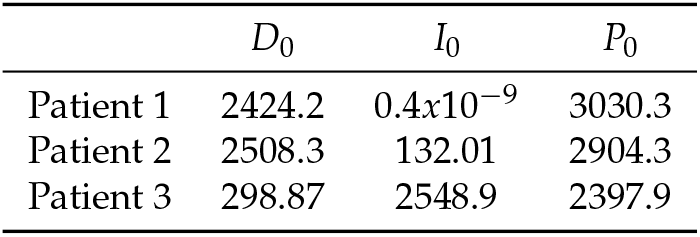
Initial conditions for different patient types

| | $D_0$ | $I_0$ | $P_0$ |
| --- | --- | --- | --- |
| Patient 1 | 2424.2 | $0.4 \times 10^{-9}$ | 3030.3 |
| Patient 2 | 2508.3 | 132.01 | 2904.3 |
| Patient 3 | 298.87 | 2548.9 | 2397.9 |

## 5. Conclusions

Adaptive therapy is a promising new approach to treating cancer. It is designed to forestall treatment resistance and prolong tumor control by competitively suppressing drug-resistant cancer cells. If its improved performance over the standard of care is due to competitive suppression, then certain design modifications should enhance its performance. Here we propose three design modifications aimed at increasing competition and improving the selection of patients who receive adaptive therapy. Using the same mathematical model that supported the original adaptive therapy design, we demonstrate that these modifications substantially improve the performance of adaptive therapy.

## Supplementary Materials

The following are available online at http://www.mdpi.com//xx/1/5/s1, Code S1: matlab code to generate Figure 3.

## Author Contributions

Conceptualization, E.H. and A.F.R.; software, E.H.; writing–original draft preparation, E.H.; writing–review and editing, E.H and A.F.R.; funding acquisition, A.F.R. All authors have read and agreed to the published version of the manuscript.

## Acknowledgments

Matlab code for simulations in Figure 3 was modeled after the code provided in the supplementary information of [13]. Landon vom Steeg, Jeanette Hansen, Valerie Morley and Don Hansen provided valuable and insightful comments on the original manuscript draft.

## Abbreviations

The following abbreviations are used in this manuscript:

ALL: Acute Lymphoblastic Leukemia
mCRPC: metastatic Castrate Resistant Prostate Cancer
PSA: Prostate Specific Antigen
ADT: Androgen Deprivation Therapy

